# Global analysis by LC-MS/MS of *N6*-methyladenosine and inosine in mRNA reveal complex incidence

**DOI:** 10.1101/2024.11.04.621854

**Authors:** Stanislav Stejskal, Veronika Rájecká, Helena Covelo-Molares, Ketty Sinigaglia, Květoslava Brožinová, Linda Kasiarova, Michaela Dohnálková, Paul Eduardo Reyes-Gutierrez, Hana Cahová, Liam P. Keegan, Mary A. O’Connell, Stepanka Vanacova

## Abstract

The precise and unambiguous detection and quantification of internal RNA modifications represents a critical step for understanding their physiological functions. The methods of direct RNA sequencing are quickly developing allowing for the precise location of internal RNA marks. This detection is however not quantitative and still presents detection limits. One of the biggest remaining challenges in the field is still the detection and quantification of m^6^A, m^6^A_m_ and m^1^A modifications. The second intriguing and timely question remaining to be addressed is the extent to which individual marks are coregulated or potentially can affect each other. Here we present a methodological approach to detect and quantify several key mRNA modifications in human total RNA and in mRNA, which is difficult to purify way from contaminating tRNA. We show that the adenosine demethylase FTO primarily targets m^6^A_m_ marks in noncoding RNAs in HEK293T cells. Surprisingly, we observe little effect of FTO or ALKBH5 depletion on the m^6^A mRNA levels. Interestingly, upregulation of ALKBH5 is accompanied by an increase in inosine level in overall mRNA.

## Introduction

The ever-growing number of RNA modifications raises the question of how to comprehensively identify different types of these modifications within a single RNA template/sample. One of the challenges is the method of choice to detect and quantify a particular RNA modification. Many methods initially relied on precipitating RNAs with a modification-specific antibody. However, the specificities of antibodies raised to different RNA modifications is highly variable (Helm *et al*, 2019). Over the past five years, there has been significant progress in establishing direct and indirect sequencing-based methods for the detection of individual RNA modifications (Dominissini *et al*, 2012; Ke *et al*, 2015; Linder *et al*, 2015; Meyer *et al*, 2012; Schaefer *et al*, 2009; Schwartz *et al*, 2014a). These approaches are excellent for qualitative detection of modifications, but are not yet so useful for quantitative assessments of multiple modifications in a given sample. To address this challenge, we employed liquid chromatography tandem mass spectrometry (LC-MS/MS), an approach that can accurately detect and quantify selected relevant modifications in a single sample.

A significant question arises regarding how the presence of one RNA modification affects the occurrence of another. As N6-methyladenosine (m^6^A) and inosine affect the same position on the adenosine and are thus mutually exclusive, it is reasonable to assume that if any RNA modification could affect the occurrence of another, it would be these two (for review see (Rengaraj *et al*, 2021)). A few reports showed negative or positive correlation between m^6^A and inosine, but the methods used were not *quantitative* (Li *et al*, 2022; Xiang *et al*, 2018). Our goal here was to globally analyze whether there were significant differences in the presence of m^6^A or inosine when their occurrence was perturbed.

The conversion of adenosine to inosine (A-to-I) in double-stranded (ds) RNA is referred to as RNA editing and is catalyzed by the adenosine deaminase acting on RNA (ADAR) enzymes (for review see (Sinigaglia *et al*, 2019)). Inosine can base pair with cytosine and is recognized primarily as guanosine during translation (Licht *et al*, 2019), potentially altering the coding sequence if present in exons. There are two active enzymes in mammals; ADAR1 and ADAR2. ADAR1 is ubiquitously expressed and is responsible for ‘promiscuous’ editing. This involves editing of duplex regions in transcripts formed by inverted repetitive elements, such as Alu elements, with an editing level of less than 1% (for review see (Eisenberg & Levanon, 2018)). Due to the high prevalence of repetitive elements in humans, there are millions of positions edited in the human transcriptome. The expression of ADAR2 in humans is high in the arteries, lungs, bladder and brain. In brain it edits specific sites of GluR transcripts by up to 100% whereas other mRNA targets show lower percentage of editing (Seeburg *et al*, 1998). We primarily focus on ADAR1 instead ADAR2 due to its wider tissue expression.

Unlike adenosine deamination, N6-methylation (m^6^A) does not change the codon meaning when it is present in an open reading frame. It is more prevalent than inosine or any other internal mRNA modification (reviewed in (Murakami & Jaffrey, 2022)). Depending on the location, m^6^A has different effects on mRNA biogenesis and function. It has been linked to alternative splicing, mRNA export, translation or mRNA stability (reviewed in (Murakami & Jaffrey, 2022)). This modification is enriched in 3’ untranslated regions (UTRs) of mRNAs and around stop codons (Dominissini *et al*., 2012; Ke *et al*., 2015), however its abundance and roles in pre-mRNAs remain for the most part unknown. Two different methyltransferases (referred to as the ‘writers’) can deposit m^6^A in mammalian mRNAs, the methyltransferase-like protein 3 and 14 heterodimer (METTL3 and METTL14) and the methyltransferase-like protein 16 (METTL16). METTL3/14 associates with additional auxiliary factors and is understood to be the main m^6^A writer responsible for majority of m^6^A in mRNAs (Liu *et al*, 2014) (Schwartz *et al*, 2014b) (Poh *et al*, 2022). METTL16 appears to have a more limited substrate repertoire. It primarily modifies *Mat2a* mRNA and U6 snRNA (Pendleton *et al*, 2017; Warda *et al*, 2017)) and few other mRNAs ((Sun *et al*, 2023; Yoshinaga *et al*, 2022) and reviewed in (Mansfield, 2024)), but it also has m^6^A independent function (Wang *et al*, 2023). In this study, we primarily address the effect of METTL3/14 on the overall m^6^A levels.

M^6^A in mRNAs can be demethylated by at least two demethylases, the so called ‘erasers’; the Fat mass and obesity-associated protein (FTO) and/or the AlkB homolog 5 RNA demethylase (ALKBH5) (Jia *et al*, 2011; Zheng *et al*, 2013). FTO displays a broader substrate specificity as it can also target m^6^A in snRNAs, *N6*, 2′-O-dimethyladenosine (m^6^Am) in mRNAs and snRNAs, and m^1^A in tRNAs (Mauer *et al*, 2019; Wei *et al*, 2018). The extent of the N6-methyl group removal ‘by the erasers’ is still controversial (Darnell *et al*, 2018), motivating us to investigate the potential dynamics of m^6^A and m^6^A_m_ modifications in detail.

Although LC-MS/MS is considered the ‘gold standard’ for the identification of RNA modifications, there are certain caveats to this method that have first to be addressed. These mainly concern the purity of the mRNA samples (Richter *et al*, 2021) and the method used to calculate the abundance of modifications. Studies that focus on mRNA require thorough sample purification. However, even after several rounds of poly(A) enrichment, “contaminating” non-coding RNAs such as rRNA can still be detected (Legrand *et al*, 2017). To address this concern, we measure tRNA- and rRNA-specific or -enriched modifications in parallel with the mRNA sample being analyzed to document mRNA purity in individual samples. We present methods for quantifying individual marks that avoid issues with the high abundance of adenosine representation in poly(A) enriched samples. Additionally, we also provide strategies for distinguishing and quantifying 5’cap-linked and internal modifications. Using these methods we show that m^6^A and RNA editing are interlinked in certain human cell types and uncover a minor role of FTO and ALKBH5 on modulating total m^6^A levels in steady-state mRNAs.

## Results

### Establishment of LC-MS/MS analysis of RNA modifications

For the RNA sample preparation and LC-MS/MS analysis we followed published protocols with some modifications (Thuring *et al*, 2016) (see methods for details). Isotope-labelled internal standards (IS) were used to determine the amount of RNA modification. These standards were prepared from total RNA isolated from *S. cerevisiae* or *E. coli* cultivated in ^13^C-labelled media. As m^6^Am is not present in bacteria or yeast, it was chemically synthesized as described in (Kellner *et al*, 2014).

We optimized the quantification of modification in both total and poly(A)-enriched RNA samples isolated from T-Rex FlpIN HEK293 cells (293T). To distinguish and quantify internal and cap-associated modifications, we performed mass spectrometry analyses of RNA samples prepared in parallel with two different nuclease treatments. Nuclease P1 (NP1) cleavage of total RNA results in digestion of all internal nucleotides to mononucleotides, while the first nucleotide remains linked to the cap (Mauer *et al*., 2019). This enables the measurement of modifications arising specifically from internal positions, which we term as ‘internal’ throughout the text (Figure 1A). To measure both internal and m^7^G cap-linked modifications (termed ‘cap+internal), we used the combination of NP1 and snake venom phosphodiesterase I (SVP), which hydrolyzes nucleotide polyphosphates (Figure 1A, Figure EV1A). Cap-linked modifications were identified through treatment with both SVP and NP1 of total RNAs, revealing the presence of m^6^Am and m^227^G, corresponding to trimethylated extended caps of snRNAs (Figure 1B). The poly(A) RNA samples showed as the most prominent marks m^7^G, m^6^Am and Am (Figure 1C).

**Figure 1.**
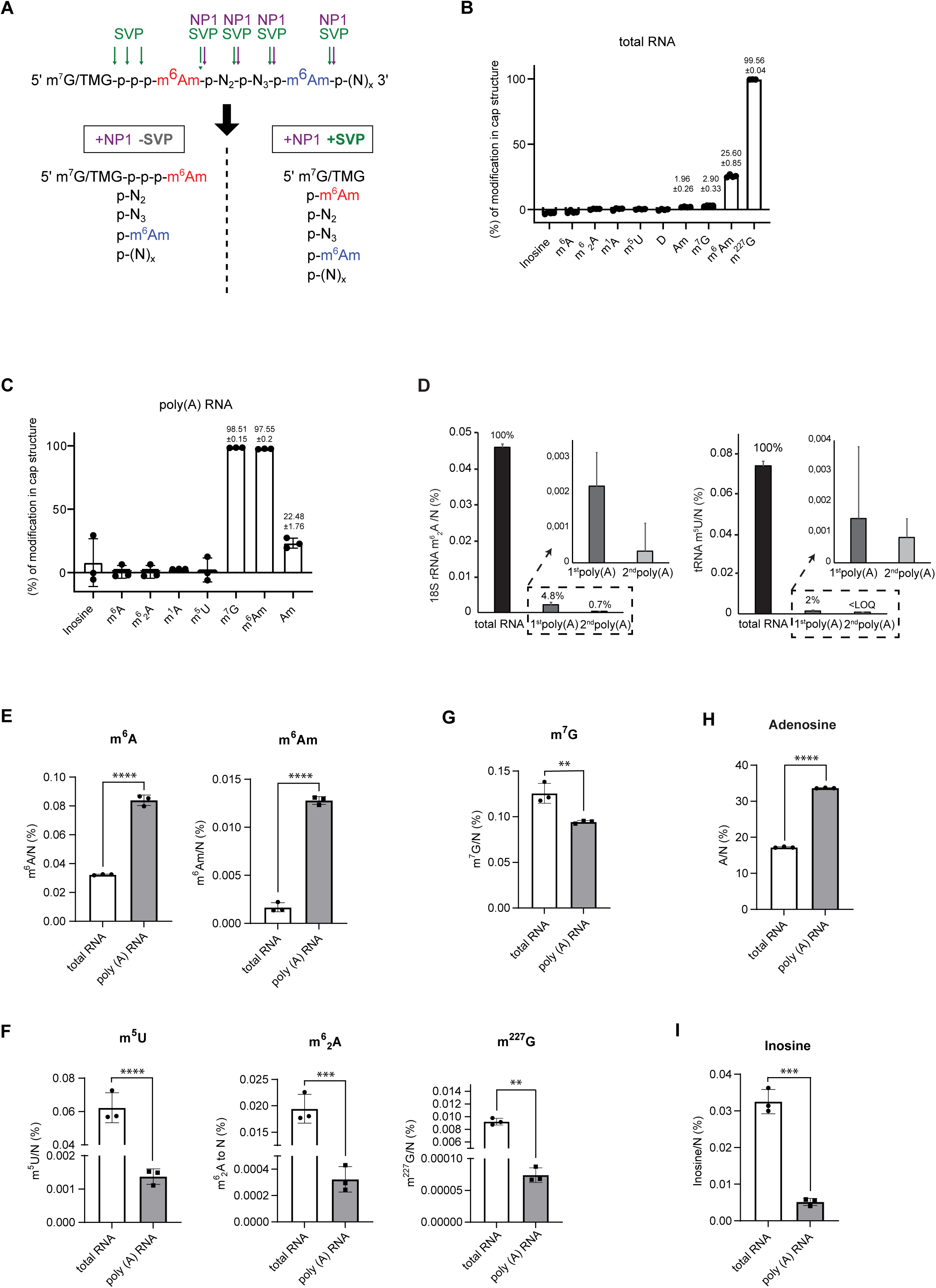
Quantitative LC-MS/MS measurements of 5’ terminal and internal modifications in total and poly(A) RNA fraction. **A** Schematic diagram illustrating the strategy for the detection of cap+internal (+NP1 +SVP) and internal (+NP1 -SVP) m^6^A_m_ levels. NP1 cleaves after each nucleotide, releasing 5’ monophosphorylated nucleotides (pN), but is unable to cleave the triphosphate bond between the RNA cap (m^7^G or m^227^G(TMG) for mRNA and snRNA, respectively) and the first nucleotide. SVP hydrolyses nucleotide polyphosphates and can therefore digest the triphosphate bond, releasing cap-linked m^6^A_m_. The green and red arrows indicate the cleavage sites of SVP and NP1, respectively. Cap-linked m^6^A_m_ in green, internal m^6^A_m_ in red. **B** Nucleoside modification content in the cap structure of total RNA in HEK293T cells (n = 3). **C** Nucleoside modification content in the cap structure of poly(A)RNA after two steps of poly(A) selection in HEK293T cells (n = 3). **D** Quantification of m^6^_2_A and m^5^U ratios in total RNA and poly(A) RNA enriched by one or two steps of poly(A) RNA isolations. The levels of nucleoside modifications were normalized to the total molar amount of canonical nucleosides N (N = C+U+G+A). n=10. Data are average ± SD. <LOQ = under the limit of quantification. **E** Measurements of m^6^A and m^6^A_m._ **F** Relative levels of m^5^U, m^6^_2_A and m^227^G, marks specific for tRNA, 18S rRNA and mature sm-class of snRNAs, respectively. Relative levels of m^7^G (**G)**, adenosine (**H**), and inosine (**I**). The levels of nucleoside modifications were normalized to the total molar amount of canonical nucleosides N (N = C+U+G+A). n=3, Data are mean ± SD. Source data are available online for this figure.

From the outset we presumed that achieving 100% purity of mRNA is not possible as it only constitutes ∼2% of cellular RNA. The presence of noncoding RNAs was thus determined by monitoring 5-methyluridine (m^5^U), which is common to tRNA and rRNA (Powell & Minczuk, 2020), *N6,N6*-dimethyladenosine (m^6^_2_A) levels as an 18S rRNA specific marker (Kellner *et al*., 2014) and *N2,N2*,7-trimethylguanosine (m^227^G) cap as a mark for PolII snRNAs and snoRNAs (Table EV1) (Buemi *et al*, 2022; Franke *et al*, 2008; Saponara & Enger, 1969; Terns & Dahlberg, 1994; Wurth *et al*, 2014). After optimization of various reagents and protocols, we achieved the best results by performing two rounds of polyadenylated RNA selection using two different poly(A) purification kits (Figure 1D) (see Methods for details). Two rounds of poly(A) selection typically resulted in significant enrichment of the abundant mRNA marks m^6^A and m^6^Am (Figure 1E), and significant depletion of the tRNA, rRNA and snRNA modifications m^5^U, m^6^_2_A and m^227^G, respectively (Figure 1F). The levels of m^7^G, a hallmark of capped mRNAs that can also be found in other PolII-derived ncRNAs, were reduced upon two rounds of poly(A) enrichment, showing depletion of ncRNAs (Figure 1G). We used the levels of canonical nucleotides for normalizations because mRNA enrichment led to a ∼20% increase in total adenosine levels and thus calculating the ratio of mRNA modifications (e.g. inosine or m^6^A) to adenosine would lead to underestimating values (Figure 1H). Compared to m^6^A, inosine levels in mRNA were by ∼6 order lower than in total RNA (Figure 1I), which is consistent with the previous data of high m^6^A prevalence in mRNAs and high prevalence of inosine in tRNAs (Figure 1I and (Torres *et al*, 2015). In summary, these results demonstrated the effectiveness of our approach to selectively enrich polyadenylated RNAs and to sufficiently deplete noncoding RNAs to be able to determine specific mRNA modifications.

### ADAR1 regulates METTL3 in a cell-type specific manner

Next, we aimed to address whether there is any cross-regulation between adenosine deamination and N6-adenosine methylation. To this end, we used HEK293T and 293T cells to determine the levels of both modifications in mRNA isolated from cells depleted of either ADAR1 or METTL3 (Figure 2A,B, Figure EV2A,B). As expected, HEK293T cells expressing shRNA targeting ADAR1 lead to a significant decrease in inosine and METTL3 knock down (KD) by siRNAs in 293T cells resulted in the reduction of m^6^A levels (Figure 2C,D). ADAR1 depletion did not alter the levels of m^6^A nor other measured marks (Figure 2C, Figures EV2A,C). Similarly, knock down of METTL3 did not result in any changes in inosine or other examined modifications (Figure 2D, Figures EV2 B,D). We did not observe any contamination of the poly(A) RNA with ncRNAs that could potentially affect the m^6^A and inosine levels (Figures EV2 C,D).

**Figure 2.**
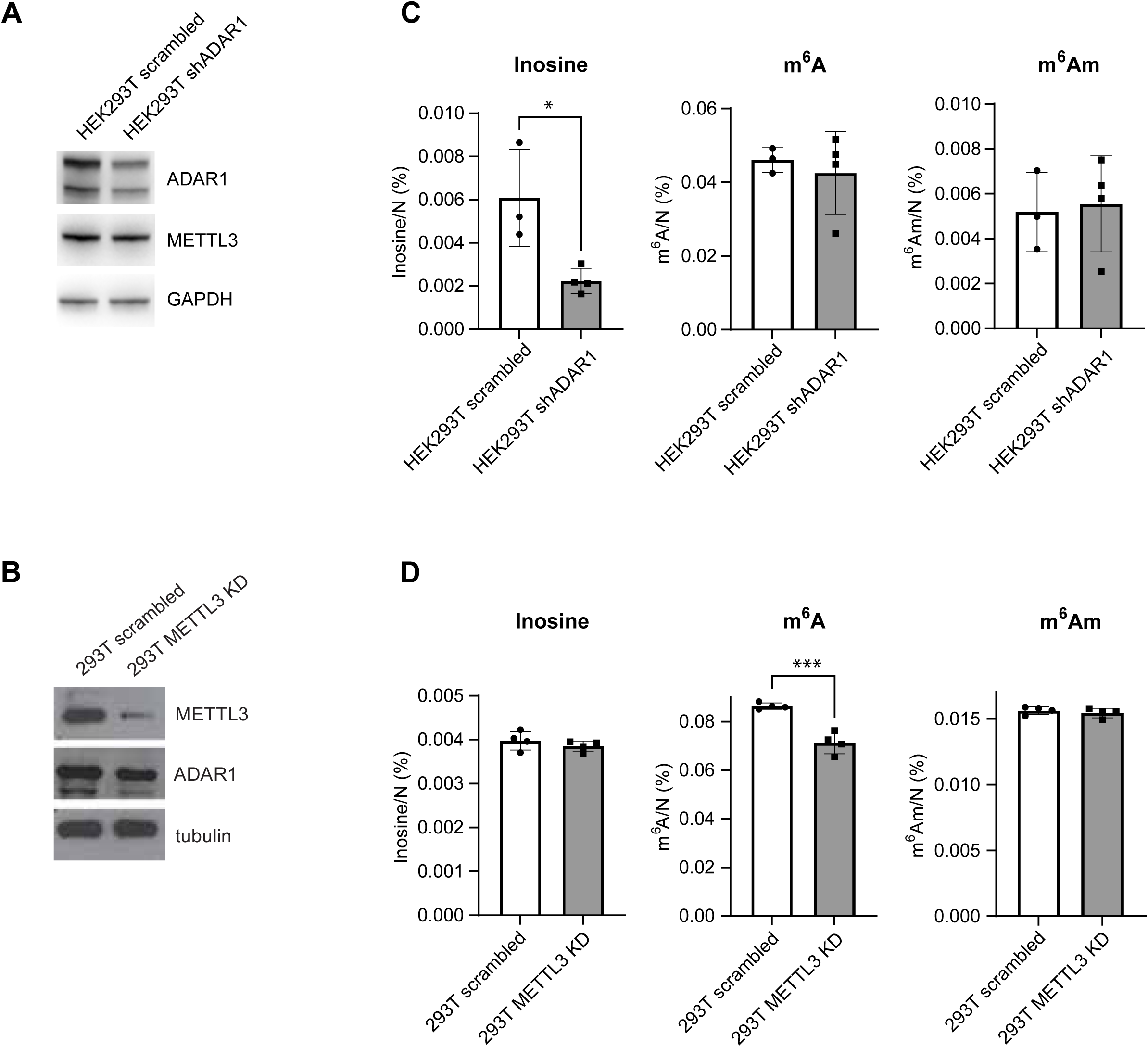
ADAR1 and METTL 3 show cross-regulation in a cell type specific manner. **A.** Immunoblot analysis of protein expression in HEK293T cells treated with control (scrambled) or shRNA specific for *ADAR1* mRNA. The levels of proteins were detected with specific antibodies listed in Materials and Methods. **B.** Immunoblot analysis of protein expression in 293T cells transfected with control (scrambled) or *METTL3* targeting siRNAs (METTL3 KD). Proteins were detected with specific antibodies listed in Material and Methods. **C** Levels of inosine, m^6^A, and m^6^A_m_ in poly(A) RNA after knockdown of ADAR1 in HEK293T cell line (n = 3-4). **D.** levels of inosine, m^6^A, and m^6^A_m_ in poly(A) RNA after knockdown of METTL3 in the 293T cell line (n = 4). Data are mean ±SD, unpaired two-tailed T-test. *P < 0.05, **P < 0.01, *** P < 0.001. The levels of nucleoside modifications were normalized to the total molar amount of canonical nucleosides N (N = C+U+G+A). Source data are available online for this figure.

### The m^6^A_m_ in ncRNAs is the primary target of FTO

In order to elucidate the dynamic potential of two common RNA modifications, m^6^A and m^6^A_m_, two knock-out (KO) cell lines of ALKBH5 KO or FTO KO were generated in 293T cells (Figure 3A). Both FTO and ALKBH5 were previously identified as m^6^A demethylases, but subsequent studies have shown that FTO exhibited a preference for m^6^A_m_ (Mauer *et al*, 2017; Wei *et al*., 2018). We initially focused on the functions of the demethylases on total RNA. As anticipated, we observed a significant increase in m^6^A_m_ levels in FTO KO cells, whereas the levels of m^6^A_m_ remained unchanged in ALKBH5 KO cells (Figure 3B). Levels of m^6^A and m^6^A_m_ methyltransferases remained unaffected in both cell lines (Figure EV3A). The observed increase in m^6^A_m_ levels in FTO KO cells was evident only upon SVP treatment, indicating that cap-linked m^6^A_m_ is a primary target of FTO (Figure 3B). In contrast, internal m^6^A_m_ levels remained unaffected in FTO KO cells (Figure 3B). This is consistent with previously published data by Mauer et al. on the demethylation role of FTO on cap-linked m^6^A_m_ in snRNAs (Mauer *et al*., 2019). Importantly, episomal expression of FTO WT, but not catalytically inactive (HD mutant) or disease-associated (RQ mutant, (Boissel *et al*, 2009)) FTO, was able to restore m^6^A_m_ levels back down to those observed in WT cells (Figure 3C). The m^6^A levels in total RNA remained unaltered following the knockout of either of the demethylases (Figure EV3B).

**Figure 3.**
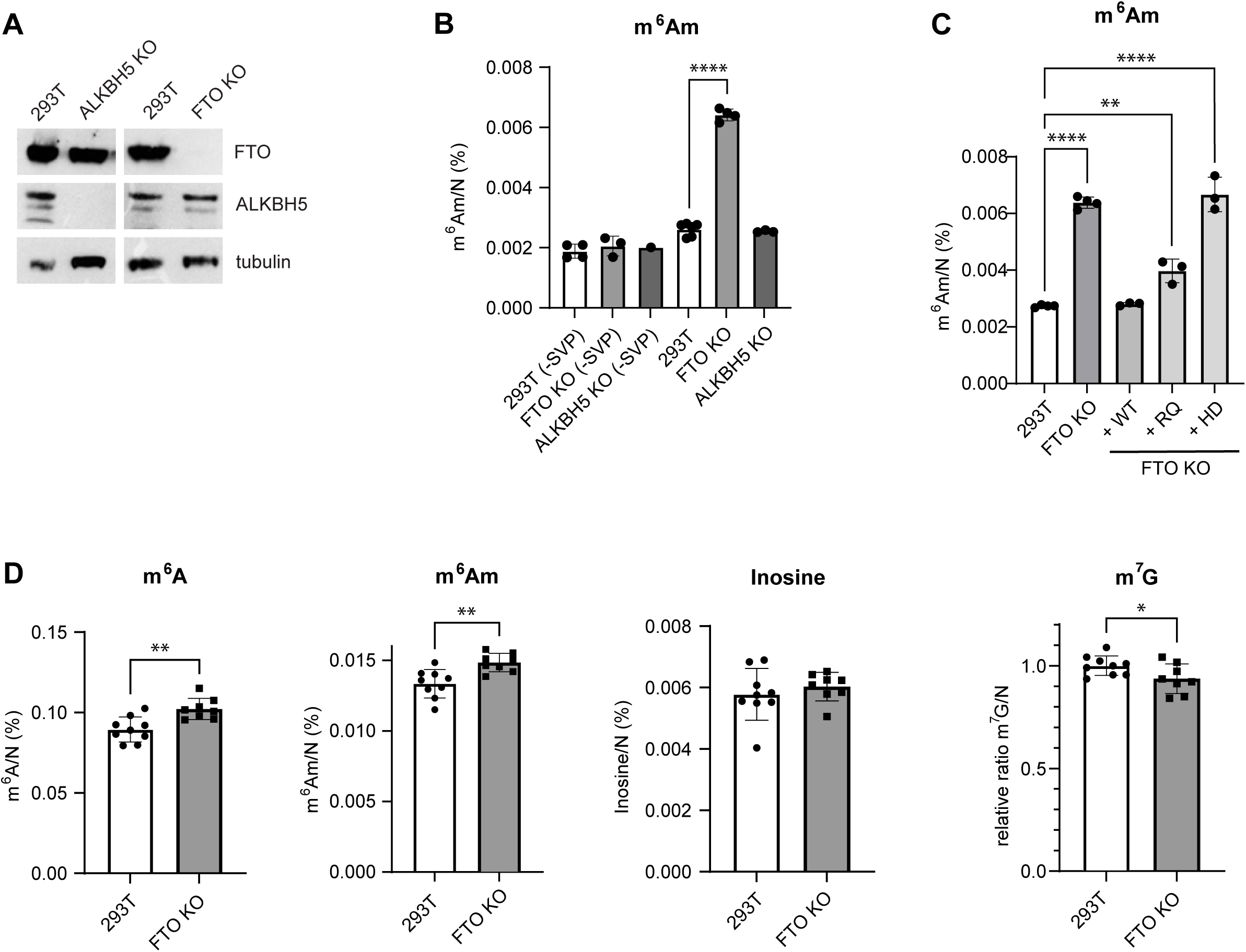
FTO primarily targets cap-associated m^6^A_m_ in noncoding RNAs. **A.** Immunoblot analysis of protein expression in 293T ALKBH5 KO and FTO KO cells. The levels of proteins were detected with specific antibodies listed in Materials and Methods. **B.** LC-MS/MS measurements and quantification of relative levels of internal (-SVP) and cap+internal m^6^A_m_ in total RNA isolated from WT, FTO KO and ALKBH5 KO 293T cells. 1 ug of total RNA was digested with NP1 and with or without SVP, to release or not the cap-linked nucleotide, respectively. In both cases, dephosphorylation with shrimp alkaline phosphatase (SAP) was performed prior to LC-MS/MS analysis. (n=3) **C.** Relative levels of m^6^A_m_ in total RNA digested with NP1 and SVP, isolated from WT, FTO KO and FTO KO cells with stably integrated forms of FTO WT (wild type), RQ (disease-linked mutant) and HD (catalytically inactive) (n=3). **D.** LC-MS/MS measurements of m^6^A, m^6^A_m_, inosine and m^7^G in poly(A) RNA isolated from WT and FTO KO cells. (n=8-9). Data are mean ± SD, One-way ANOVA, multiple comparison using the Tukey test. *P < 0.05, **P < 0.01, *** P < 0.001. The levels of nucleoside modifications were normalized to the total molar amount of canonical nucleosides N (N = C+U+G+A). Source data are available online for this figure.

To further elucidate the dynamics of m^6^A(m) upon dysregulation of FTO we analyzed poly(A) RNA from FTO KO cells. We observed a small but significant increase in both m^6^A and m^6^A_m_ levels in FTO KO compared to WT 293T cells when combining nine independent measurements (Figure 3D), which is consistent with a previous report (Wei *et al*., 2018). This m^6^A and m^6^A_m_ stabilization in mRNAs had no impact on the level of inosine in mRNAs (Figure 3D). We observed a slight reduction in m^7^G levels in FTO KO cells (Figure 3D) but this could be due to slightly higher tRNA or rRNA contamination in the WT poly(A) RNA samples, as indicated by m^5^U levels (Figure EV3C), even though this contamination did not affect the levels of m^6^A (Figure 3D). In summary, FTO primarily targets cap-linked m^6^A_m_ in total RNA, but also has a relatively minor yet significant effect on m^6^A and m^6^A_m_ levels in poly(A) RNA of 293T cells. These results suggest that m^6^A_m_ modification displays a dynamic potential in steady-state levels of total RNA, while the effects on poly(A) RNA are rather small in this cell line.

### ALKBH5 depletion has little impact on m^6^A levels in steady state mRNAs

ALKBH5, a conserved m^6^A RNA demethylase found in most eukaryotes, is considered to be the main eraser of m^6^A in mRNA. To date, its activity has been linked to protein-coding transcripts (reviewed in (Rajecka *et al*, 2019)). Given that FTO depletion revealed little impact on mRNA m^6^A stabilization, we next investigated how ALKBH5 dysregulation affects m^6^A and other mRNA modifications. Firstly, we compared m^6^A levels in poly(A) RNA in control 293T cells and ALKBH5 KO cells. Surprisingly, we observed only a very small and insignificant increase in m^6^A (Figure 4A) and no changes in the other marks evaluated when comparing ALKBH5 KO cells with WT HEK293T (Figure 4A, Figure EV4A). We hypothesized that the CRISPR-generated ALKBH5 KO cells might have adapted their RNA metabolism to the lack of ALKBH5. Therefore, we performed conditional ALKBH5 depletion by siRNAs and extended the study to several other human cell lines apart from 293T cells (Figure 4B). Similarly to FTO depletion, we detected only a very mild increase in m^6^A in poly(A) RNA in 293T and HeLa cells and no significant change for the other two tested cell lines (Figure 4C). ALKBH5 downregulation did not affect levels of any of the other measured modifications (Figure 4C, Figure EV4C) Interestingly, the relative levels of poly(A) RNA m^6^A, m^6^A_m_ , inosine and m^7^G modifications differed between individual cell lines and these changes do not seem to be explained by differences on writer’s expression between cell lines (Figure 4C, Figure EV4B,C). On contrary, overexpression of ALKBH5 (Figure 4D) led to a strong and significant decrease in m^6^A and small but significant increase in inosine in the poly(A) RNA fractions (Figure 4E). The other determined RNA modifications as well as protein level of methyltransferases remained unaffected (Figure EV4D, EV4E).

**Figure 4.**
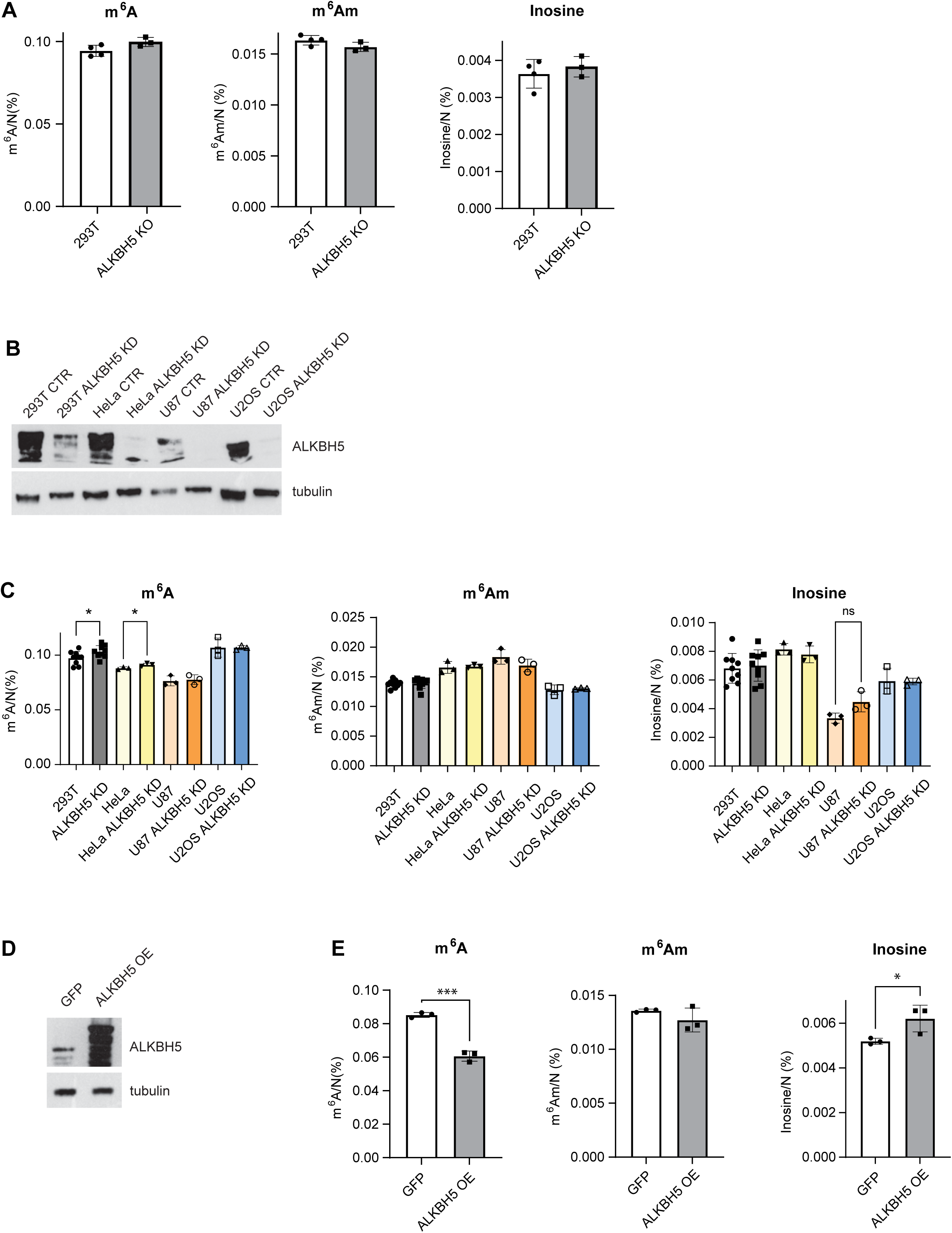
ALKBH5 depletion has a minor effect on m^6^A levels in poly(A) RNA. **A.** LC-MS/MS measurements of m^6^A, m^6^A_m_ and inosine in poly(A) RNAs isolated from WT and ALKBH5 KO cells. **B.** Immunoblot analysis of protein expression in 293T, HeLa, U87 and U2OS cells transfected with control (scrambled) or *ALKBH5* targeting siRNAs (ALKBH5 KD). The levels of proteins were detected with specific antibodies listed in Materials and Methods. **C.** LC-MS/MS measurements of m^6^A, m^6^A_m_ and inosine in poly(A) RNAs isolated from WT and ALKBH5 KD in 293T, HeLa, U87 and U2OS cells. 293T (n=9), HeLa, U87 and U2OS (n=3). **D.** Immunoblot analysis of protein expression in 293T cells overexpressing ALKBH5 (ALKBH5 OE) or GFP control. The levels of proteins were detected with specific antibodies listed in Materials and Methods. **E.** LC-MS/MS measurements of m^6^A, m^6^A_m_ and inosine in poly(A) RNAs isolated from control (GFP) and ALKBH5 overexpressing cells. Data are mean ± SD, One-way ANOVA, multiple comparison with the Tukey test (for ALKBH5 KD). Data are mean ±SD, unpaired two-tailed T-test (for ALKBH5 KO and ALKBH5 OE). *P < 0.05, **P < 0.01, *** P < 0.001. The levels of nucleoside modifications were normalized to the total molar amount of canonical nucleosides N (N = C+U+G+A). Source data are available online for this figure.

In summary, ALKBH5 depletion demonstrated minimal to no effect, exerting a nearly zero effect on m^6^A levels in a steady-state pool of mRNAs, with notable cell-type-dependent variations. Only strong upregulation of ALKBH5 shows a strong m^6^A eraser effect.

## Discussion

The aim of this study was to investigate the dynamics of m^6^A, m^6^A_m_ and inosine and to examine whether levels of m^6^A and inosine, the two RNA modifications that are amongst the most prevalent in mRNA, influence each other. Both of these modifications target the same position on the adenosine, so they are mutually exclusive. However, they occupy distinct sequence-specific positions and RNA structures, precluding direct exchange of the modifications at the same position. Consequently, any potential changes in m^6^A or inosine levels would be indirect. We chose LC-MS/MS to perform the analysis as we want to observe the global levels of m^6^A, m^6^A_m_ and inosine upon dysregulation of the enzymes involved in generating or removing these modifications. Analyzing different modifications in parallel in the same samples should mean that any artifact introduced by this method would have an effect on both modifications, limiting the potential biases of the analysis.

In our study we initially focused on the establishment of an LC-MS/MS method that would allow us to measure cap-linked and internal modifications. Previous studies have quantified cap-linked dinucleotides upon NP1 RNA digestion (Galloway *et al*, 2020; Mauer *et al*., 2019; Wang *et al*, 2019) or used NP1 together with a pyrophosphatase for cleavage to single nucleotides (Boulias *et al*, 2019; Sendinc *et al*, 2019). We established a method that allowed us to distinguish between internal and cap-linked modifications by combining the NP1 and SVP digestion. The measurement of RNAs digested by NP1 alone or in combination with SVP enabled the identification of cap-linked modifications. Secondly, previous mass spectrometry studies on m^6^A in poly(A) RNA did not address the potential significant contamination resulting from the ineffective isolation of poly(A) RNA from total RNA (Wei *et al*., 2018; Zheng *et al*., 2013). Consequently, we focused on the measurement of tRNA and rRNA-specific modifications, which would allow us to monitor the levels of tRNA and rRNA contamination in each sample. By documenting the purity of the poly(A) RNA samples we can reflect m^6^A levels independently of tRNA or rRNA contamination.

Our results of the m^6^A-inosine interconnection studies suggest that the responsible proteins, METTL3 and ADAR1, might affect the levels of the cross-related marks only in a cell-type dependent manner. The knockdowns of either METTL3 or ADAR1 in 293T or HEK293 cells, respectively, did not lead to changes in the other modifications. These observations are consistent with the latest reports on the cross-regulation between METTL3 and ADAR1. Terajima et al. demonstrated that the downregulation of METTL3 does not affect ADAR1 levels in the glioblastoma A172 cell line, unless the interferon response is stimulated (Terajima *et al*, 2021). HEK293T cells do have a strong interferon response so it is not certain that adding interferon would change the level of m^6^A. In contrast, Li et al. observed that ADAR1 activity positively regulates METTL3 expression in breast cancer cells (MCF7 and MDA-MB-231). In that case, ADAR1 modifies the miR-532-5p seed sequence in METTL3 mRNA, leading to its stabilization, enhanced translation and consequently to higher general m^6^A levels in mRNAs (Li *et al*., 2022). It is currently unclear why a similar ADAR1-mediated regulation of METTL3 was not observed in HEK293 cells. The miR532-5p is relatively highly expressed in HEK293T cells (miRmine). ADAR1 has many interacting proteins and some are differentially expressed in different cell lines (Vukic *et al*., accepted for publication in NAR). Thus, it may employ different mechanisms to regulate METTL3 expression in different cell types. Our findings, when considered alongside the existing published data, indicate that the consequence of METTL3 editing is cell-type dependent at the transcriptome-wide level. Furthermore, we did not observe any clear cross-regulation between m^6^A, m^6^A_m_ and ADAR1.

We further focused on the investigation of m^6^A_m_ as a reversible mark. The double digestion of RNA with NP1 and SVP was employed to quantify m^6^A_m_ levels in RNA isolated from FTO KO cells. We detected the previously reported changes in m^6^A_m_ abundance in total RNA samples (Mauer *et al*., 2019) and confirmed previously published data indicating that m^6^A_m_ in total RNA is a primary target of FTO (Mauer *et al*., 2017). M^6^A_m_ levels in total RNA in FTO KO cells could be restored by reintroduction of WT FTO, but not by the introduction of catalytically inactive HD mutant FTO (in α-ketoglutarate coordination site H231A, D233A). The disease-associated RQ mutant FTO (in the 2-oxoglutarate coordination site R316Q, (Boissel *et al*., 2009)) showed an m^6^A_m_ abundance between that observed in 293T WT and HD mutant FTO cells. This is particularly noteworthy because our previous findings indicated that RQ mutant cells exhibit different alternative splicing patterns in specific transcripts between those observed in WT and HD mutant FTO. This suggested that the splicing changes that we observed in FTO KO 293T cells (Bartosovic *et al*, 2017) may be regulated by m^6^A_m_ levels in snRNAs. The levels of m^6^A_m_ in poly(A) RNA were significantly increased upon FTO KO, in agreement with previously published data (Wei *et al*., 2018). Taken together, our findings indicate that m^6^A_m_ exhibits reversibility and reveals dynamic potential in both total RNA and poly(A) RNA.

In regard to the measurements of m^6^A, our data do not support the prevailing notion that m^6^A is a dynamic modification. Our findings are consistent with those of the Darnell and Jaffrey groups (Darnell *et al*., 2018; Meyer & Jaffrey, 2017), who posit that m^6^A is a stable modification. Firstly, m^6^A levels remained almost unaffected in FTO-dysregulated cells in both total RNA and poly(A) RNA, supporting the hypothesis that FTO targets m^6^A in mRNAs only minimally (Mauer *et al*., 2017) and suggesting that m^6^A is rather more a stable than a dynamic RNA modification. Secondly, the depletion of ALKBH5 showed only marginal effect on m^6^A levels in only some cell lines that we analyzed. In particular, in order to achieve statistical significance for a rather small overall stabilization of m^6^A, we had to perform a higher number of measurements than the minimum of three replicates usually required for statistical tests. Therefore, the observed results are statistically significant, although their biological relevance is rather unclear. Our observations on the ALKBH5 demethylase activity align with those reported by other groups using LC-MS/MS measurements (Zheng *et al*., 2013). Only high overexpression of the eraser led to a dramatic decrease of m^6^A in poly(A) RNA, as observed in the aforementioned study (Zheng *et al*., 2013). These data are consistent with the findings of other reports on cancers that have demonstrated a correlation between elevated ALKBH5 expression in various cancer types and decreased m^6^A levels (Chao *et al*, 2020; Jiang *et al*, 2020; Zhang *et al*, 2016). The elevated levels of ALKBH5 have been identified as an oncogene that can facilitate the growth of cancer cells (Jin *et al*, 2022; Qu *et al*, 2022a). However, several studies have also demonstrated its tumour suppressor function (reviewed in (Qu *et al*, 2022b)), thereby revealing its dual role in carcinogenesis. Furthermore, our findings challenge the prevailing view that ALKBH5 is primarily an m^6^A demethylase, as its demethylase activity has been predominantly reported when it is upregulated in cancer, although several studies have also demonstrated its demethylase function also in non-cancer research (Gao *et al*, 2023; Yu *et al*, 2020). ALKBH5 KO cells and animals are viable (Bai *et al*, 2023; Gao *et al*., 2023). ALKBH5 KO mice show defects in spermatogenesis and oogenesis (Bai *et al*., 2023; Tang *et al*, 2018). This is in contrast to the original proposition that functional m^6^A dynamicity is essential for basic cellular functions (Jia *et al*., 2011; Meyer & Jaffrey, 2014). ALKBH5 is capable of binding the m^6^A through its m^6^A-binding pocket (Aik *et al*, 2014; Feng *et al*, 2014), thus it remains to be determined whether ALKBH5 could act as an m^6^A reader rather than a demethylase.

In conclusion, the data presented suggest that m^6^A_m_ is a potential dynamic reversible RNA methylation mark in both pools of steady-state total RNA (presumably snRNAs) and poly(A) RNA. However, further experiments are required to provide more definitive evidence. In contrast, m^6^A, which was previously believed to be a dynamic mark, demonstrated close to no reversible potential *in vivo* on overall steady-state poly(A) RNAs. Many studies pointed to specific roles of m^6^A in splicing, stability or translation of single particular mRNAs that often occur cell and/or tissue-specifically (He & He, 2021; Murakami & Jaffrey, 2022). Overall, our evidence does not support the hypothesis that epitranscriptomic RNA modification in general should be dynamic and resemble epigenetic modifications.

## Materials and Methods

### Human cell line manipulation

Human 293 Flp-In™ T-REx™ (293T) (Invitrogen) cells were maintained as described in (Covelo-Molares *et al*, 2021). Human HEK293T (ATCC, CRL-3216), T-REx-HeLa cells (Invitrogen), U87 and U2OS cells were cultured in Dulbecco’s Modified Eagle Medium (DMEM) supplemented with 10% fetal bovine serum (FBS) at 37°C in the presence of 5% CO2.

### Preparation of FTO and ALKBH5 knock-out (KO) cell lines by CRISPR-Cas9

To create cells with disrupted expression of FTO or ALKBH5, we employed the CRISPR/Cas9 system (Ran *et al*, 2013). We used the following sgRNAs targeting intron 2 and exon 3 of the *FTO* gene: FTO-intron2: CACCgACTCGTGCCCTGGAAGCCAG; FTO-exon3: CACCgTATGTCTGCAGATTTCCCCA, and the following sgRNAs targeting exon 1 of the *ALKBH5* gene: ALKBH5-exon1a: CACCgACGTCCCGGGACAACTATA; ALKBH5-exon1b: CACCgTGGACTTGAGCTTCTCACGC. The oligonucleotides were individually cloned into the vectors pSpCas9(BB)-2A-GFP (PX458) (from Feng Zhang, Addgene #48138) for FTO and pSpCas9n(BB)-2A-Puro (PX462) V2.0 (from Feng Zhang, Addgene #62987) for ALKBH5, and verified by Sanger sequencing. On the day of transfection, 250ng of each sgRNA construct was mixed with serum-free DMEM and transfected with the Lipofectamine3000 reagent (Invitrogen, L3000015) to 50% confluent cells. For FTO KO cells, 24 hours post-transfection cells were single-cell sorted into a 96-well plate with FACS. For ALKBH5 KO, 24 hours post-transfection puromycin (Applichem, A2856) was added (final concentration 4ug/ml) and cells were grown to 60% confluency. A second round of transfection was performed as described above. 24 hours post-transfection, cells were counted and seeded at the density of 0.5 cell/well in 96-well plates.

The individual cells were verified as KO by immunoblot, RT-qPCR, and Sanger sequencing of the corresponding genomic DNA locus. Expression of proteins was analysed in selected clones by immunoblot. *FTO* and *ALKBH5* mutations/deletions were validated by PCR on the genomic DNA with primers FTO_Fwd: 5’TCATTCATCCATGCACAAATCC3’, FTO_Rev: <colcnt=4> 5’GCAGAGCAGCATACAACGTA3’, ALKBH5_Fwd: 5’ CCGTTGTCGCCACCGTTGCATGAC3’ and ALKBH5_Rev: 5’ AGTCCTCCTGATACTTGCGCTTGG 3’.

### RNA isolation

Total RNA was isolated with TriPure reagent (Sigma, T3934-100) according to the manufacturer’s protocol followed by a precipitation step with 1 volume of 5M ammonium acetate (Invitrogen, AM9070G) and 2.5 volumes of 100% ethanol. RNA integrity was assessed by electrophoresing an aliquot of the RNA sample on a denaturing agarose gel stained with SYBR® safe (Invitrogen, S33102).

### Poly(A) mRNA enrichment

1mg of total RNA was used as input for the first step of poly(A) mRNA enrichment with the PolyATtract® mRNA Isolation Systems kit (Promega, Z5310) and purified according to the manufacturer’s protocol. Eluted poly(A)+ mRNA was precipitated with 1 volume of 5M ammonium acetate (Life Technologies) and 2.5 volumes of 100% ethanol. After mRNA recovery, the second step of poly(A) mRNA enrichment was performed with Poly(A) RNA Selection kit (Lexogen, 039). To increase the yield of mRNA, 3x the suggested amount of oligo dT magnetic beads were used. The concentration of eluted mRNA was measured by Qubit^TM^ RNA HS Assay kit (Invitrogen, Q32852). Typically, 500ng-2ug of mRNA was obtained after the two rounds of poly(A) mRNA enrichment.

### LC-MS/MS quantitative analysis of RNA modifications

The analysis of RNA modifications by LC-MS/MS was performed according to the protocol developed by Thuring and co-workers (Thuring et al., 2016). Carbon-13 (^13^C)-labelled internal standards (IS) were prepared from bacterial and yeast cultures. *Escherichia coli* strain BL21 (DE3)-RIL was cultivated in M9 minimal medium (3.37 mM Na2HPO4, 2.20 mM KH2PO4, 0.86 mM NaCl, 0.94 mM NH4Cl, 1.00 mM MgSO4, 0.3mM CaCl2) supplemented with Trace elements solution (final concentration – 134.00 µM EDTA, 31.00 µM FeCl_3_, 0.62 µM ZnCl_2_, 0.76 µM CuCl_2_, 0.42 µM CoCl_2_, 1.62 µM H_3_BO_3_, 0.08 µM MnCl_2_) and 20% D-GLUCOSE-^13^C_6_ (Merck, 389374). Bacteria were harvested at late exponential phase (OD_600_ = 1.8), centrifuged at 6000*g* 10 min 4°C and twice washed with cold 1xPBS. RNA was isolated with TRI Reagent® (Merck, T9424) according to the manufacturer’s protocol followed by a precipitation step with 1 volume of 5M ammonium acetate and 2.5 volumes of 100% ethanol.

*Saccharomyces cerevisiae* strain BY4741 was grown in ^13^C-labeled rich growth yeast media (OD_600_ = 2) (SILANTES, 111201402) supplemented with 20% Glucose ^13^C_6_ (SILANTES, 302204100) at 30°C for 12-14 hours with shaking at 230-270 rpm. Yeast harvested at the late exponential phase (OD_600_ approximately 3.5), centrifuged at 3500*g* 10 min 4°C and washed two times with ice-cold 1xPBS. The cell pellet was resuspended in 1 ml of TRI Reagent® per 10 ml of SILANTES media. The supernatant was transferred to pre-fill screw-capped vials with ∼250 μl volume of acid-washed beads (500 µm diameter, BIOSPEC). Cells walls were disrupted with Precellys evolution homogenizer (Bertin Technologies) 4x (2.5 min homogenization at 6500 RPM/ 2.5 min sample rest on ice). Debris from the transferred suspension was removed by centrifugation at 6000*g* 15 min 4°C and the supernatant was used for RNA isolation as previously described.

### Synthesis of *N*^6^-(methyl-d_3_)-2’-*O*-methyladenosine (m6A_m_-d_3_) standard

A mixture of 2′-*O*-methyl-adenosine (250 mg, 0.89 mmol) and methyl iodide-d_3_ (221 ul, 3.5 mmol) in 2.0 mL of *N*,*N*-dimethylacetamide (DMA) was stirred at 28°C overnight. The reaction mixture was then treated with 20 mg of celite and stirred for 10 min. The suspension was filtered and acetone (10 mL) was added to wash the solid part which was discarded. Crude product was precipitated from the liquid by addition of Et_2_O (10 ml). The suspension was sonicated, centrifuged, and liquid was separated from the solid pellet. To purify the precipitate, MeOH (1.0 ml) and Et_2_O (40 ml) were added to the solid pellet. The resulting suspension was sonicated, centrifuged, and the liquid was separated from the solid pellet. This process was repeated twice. Finally, Et_2_O (10 ml) was added to the solids and the suspension was sonicated and then centrifuged. Removal of liquid, and drying of the solid product at RT (5-10 min) under vacuum (3.0 mbar) led to the production of crude *N*^1^-(methyl-d_3_)-2’-O-methyladenosine as a white off solid (350 mg).

*N*^1^-(methyl-d_3_)-2’-O-methyladenosine was dissolved in 0.25 M NaOH solution (15 ml) and heated overnight at 80°C. The reaction mixture was subsequently cooled to RT and pH was adjusted to 7.5 by addition of an aqueous 10% *p*-toluensulfonic acid solution. Water was removed under vacuum (72 mbar) at 50°C. MeOH (50 ml) was added to the resulting solids and stirred for 10 min at 50°C. MeOH was subsequently removed under vacuum (337 mbar) at 40°C. The solids were refluxed overnight in EtOAc (50 ml). The resulting suspension was cooled at RT and EtOAc was removed under vacuum (240 mbar) at 40°C. MeOH (1.0 ml) and Et_2_O (40 ml) were then added to the solids. The resulting suspension was sonicated, centrifuged, and the liquid was separated from the solid pellet. The purification process was repeated an additional four times. Finally, Et_2_O (10 ml) was added to the solids and the suspension was sonicated and centrifuged. Removal of liquid and drying of the solid product at RT (5-10min) under vacuum (3.0 mbar) led to the generation of *N*^6^-(methyl-d_3_)-2’-*O*-methyladenosine (m^6^A_m_-d_3_) as a white off solid (150 mg crude product, 0.542 mmol, 56% yield). A fraction of the crude product was purified by preparative HPLC (A-Triethylammonium acetate 0.1 M, pH 7, B-Acetonitrile). Co-distillations with water followed by several freeze-drying from water gave off-white solid product.

### RNA digestion and LC-MS/MS analysis

2 µg of total RNA or 750 ng of poly(A) mRNA (or ^13^C-labelled internal standard) was digested with 0.1U nuclease P1 (NP1, Merck, N8630) and 0.3U Snake venom phosphodiesterase (SVP, Worthington, LS003928) to detect internal and cap-linked modifications in a buffer containing 0.02 mM ZnCl_2_, 20 mM ammonium acetate, pH 5.0 for 2 hours at 37°C. The digestion mix was supplemented with adenosine deaminase inhibitor Pentostatin (0.7 µg). Digested nucleotides were dephosphorylated by adding 0.3U Shrimp alkaline phosphatase (Life Technologies) and 0.1 volume of 10x dephosphorylation buffer (100 mM MgCl_2_, 100 mM ammonium acetate, pH 9.0) for 1 hour at 37°C. Internal standards were also added to the reaction: 50 ng of bacterial IS, 150ng of yeast IS and 60pg of m^6^A_m_-d_3_.

Digested nucleosides were fractionated with YMC-Triart C18 column (100 × 3.0 mm I.D., S – 3µm, 12 nm, YMC) at 35°C with HPLC Agilent 1260 infinity (Agilent). The injection volume was 10 µl (250 ng RNA). The solvent system consisted of 0.1% formic acid/water (solvent A) and acetonitrile/0.1% formic acid (solvent B). A gradient elution program was applied at a flow rate of 0.35 ml.min^-1^ at 35°C as follows: 0 min, 100% solvent A; 0 to 15 min, (92% A, 8% B); 15 to 20 min, (60% A, 40% B); 20 to 23 min, (60% A, 40% B); subsequently a column equilibration step was applied – 23.1 to 34 min (100% A). The molecular concentration of canonical nucleosides (C, U, G, A) was measured with 1260 infinity DAD detector with InfinityLab Max-Light cartridge cell, 10 mm, 1.0 μl (Agilent). The signal was analyzed at 254 nm wavelength; 4 nm bandwidth; 360 nm reference wavelength; 100 nm reference bandwidth. Eluted nucleotides were ionized in Agilent Jet stream electrospray source with the following parameters: positive mode, gas temperature 350°C, gas flow 8l.min^-1^, Nebulizer 50 psi, Sheat gas temperature 350°C, Sheat gas flow 12l.min^-1^, capillary voltage (3000V). Nucleosides were quantified with the Agilent 6460 Triple Quad Mass Spectrometer.

### Downregulation of gene expression by siRNAs

SiRNAs were transfected into cells with Lipofectamine RNAiMAX (ThermoFisher, 13778150) according to manufacturer’s protocol. For METTL3 KD, we used two different siRNAs targeting METTL3 CDS (CUGCAAGUAUGUUCACUAUGA[dT][dT], AGGAGCCAGCCAAGAAAUCAA[dT][dT], Sigma) at concentration 40 nM (20 nM each siRNA). Cells were collected 96h after transfection. For ALKBH5 KD, four different siRNAs targeting ALKBH5 CDS (ACAAGUACUUCUUCGGCGA[dT][dT], GCGCCGUCAUCAACGACUA[dT][dT], CUGAGAACUACUGGCGCAA[dT][dT], AAGUCGGGACUGCAUAAUUAA[dT][dT], Sigma) were used at concentration 20 nM (5nM each siRNA). Cells were collected 72 hours (24h+48 hours Fwd transfection) after transfection. For ADAR1 KD, two different siRNAs targeting ADAR1 3’UTR (GAUUCUUAACUGCUACAGAUA[dT][dT], UAUCUGUAGCAGUUAAGAAUC[dT][dT]) were used at concentration 30nM (15 nM each siRNA) and collected 48 hours after transfection. For negative controls of the RNAi treatment we used scrambled siRNA pool (ON-TARGETplus Non-targeting Control Pool, Dharmacon, D-001810-10-20) and scrambled siRNA pool (MISSION® siRNA Universal Negative Control #1, Sigma, SIC001) at the same concentrations as the shRNAs targeting specific mRNAs.

### ADAR1 KD by shRNAs

A stable human embryonic kidney cell lines HEK293 expressing the SMARTvector Inducible Lentiviral shRNA against ADAR1 (Horizon Discovery) or scrambled shRNA were generated following the protocol described in (Tassinari *et al*, 2021). 1 µg/mL puromycin was used for the selection two days post infection. Vector expression was induced with 1 µg/mL doxycycline refreshed every 48 hours. GFP^+^ cells were sorted after 48 hours of induction, and they were maintained in DMEM supplemented with 10% FBS (Gibco-Life Technologies), 100 U/ml penicillin and 100 µg/ml streptomycin, at 37°C in 5% CO_2_. Cells were collected after 96 hours of doxycycline induction.

### Protein overexpression in human cell lines

For protein overexpression of ALKBH5 and GFP (control) we used stable inducible cell lines previously described in (Covelo-Molares *et al*., 2021). The cells contain stable integrated fusion versions of Strep-HA-tag-ALKBH5 and STREP-HA-tag-GFP, respectively. For protein overexpression of FTO and its catalytically inactive HD mutant (in α-ketoglutarate coordination site H231A, D233A) and the disease-associated RQ mutant (patient mutation in *FTO* in the 2-oxoglutarate coordination site R316Q (Boissel *et al*., 2009)) in FTO KO cell line, we used stable inducible cell lines previously described in (Bartosovic *et al*., 2017). The cells contain stable integrated fusion versions of 3xFLAG-FTO-WT, 3xFLAG-FTO-HD or 3xFLAG-FTO-RQ, respectively. Cells were induced with 200 ng/ml doxycycline at 50% confluency and harvested after 24 hours further incubation.

### Protein expression analysis

For immunoblot analyses of HEK293T, FTO KO, ALKBH5 KO and cells overexpressing Strep-HA-tag-GFP and Strep-HA-tag-ALKBH5, area corresponding to one 6-well was collected from 15cm plates before proceeding to RNA isolation. For immunoblot analysis of HEK293T, HeLa, U87 and U2OS ALKBH5 KD cells, an area corresponding to half of 6-well was collected from 10 cm plates before further proceeding to RNA isolation. 30 μg of total cell extract was analyzed by SDS-PAGE and immunoblot with protein-specific antibodies was performed. Antibodies were used at the following dilutions: METTL3 1:1000 (Proteintech, 15073-1-AP), METTL14 1:1000 (Sigma, HPA038002-100UL), METTL16 1:1000 (OriGene, TA504710), PCIF1 1:1000 (Cell Signaling Technology, E8B1B), FTO 1:5000 (Abcam, ab126605), ALKBH5 1:3000 (Sigma, HPA007196), ADAR1 1:3000 (Antibodies-online, ABIN2855100), ADAR2 1:500 (Santa Cruz, Sc-73409), MDA5 1:800 (Cell Signaling Technology, D74E4), RIG-I 1:800 (Cell Signaling Technology, D14G6), TUB1 1:8000 (Sigma, T6074), GAPDH 1:40000 (Proteintech, 60004-1-Ig).

## Acknowledgements

We would like to thank Karolina Vavrouskova and Leona Kledrowetzova for excellent technical support. This work was supported by the Ministry of Education, Youth and Sports of the Czech Republic grant RNA for therapy (CZ.02.01.01/00/22_008/0004575) and within programme INTER-COST (LTC18052), the Czech Science Foundation (19-21829S and 22-12871S) to SV, (19-16963S) to LK, (21-27329X) to MAO’C, the institutional support CEITEC 2020 (LQ1601). HCM was supported by Brno City Municipality Scholarship for Talented Ph.D. SS was funded by Czech Science Foundation (20-11101S). HC and PER-G acknowledge funding from the Ministry of Education, Youth and Sports (Czech Republic), programme ERC CZ (LL1603).

## Authors contribution

Stanislav Stejskal: established the method and performed analyzed RNA samples on LC-MS/MS.

Veronika Rájecká: produced results concerning FTO and ALKBH5 analyses, performed KD in cell lines, isolated total RNA and mRNA for analysis, performed immunoblot analyses and participated at data interpretation and manuscript writing Helena Covelo-Molares: initiated the project, produced results concerning FTO analyses, established the method of mRNA isolation from human cell lines, designed part of the experiments, isolated total RNA and mRNA for analysis, interpreted the results, participated in manuscript writing.

Ketty Sinigaglia: established the method, produced results concerning ADAR1 KD, performed KD in cell lines, isolated mRNA for analysis.

Květoslava Brožinová: performed analyzed RNA samples on LC-MS/MS Michaela Dohnálková: Prepared and validated the ALKBH5 KO cell line. Linda Kasiarova: Prepared and validated the FTO KO cell line.

Paul Eduardo Reyes-Gutierrez: synthesized *N*^6^-(methyl-d_3_)-2’-*O*-methyladenosine (m6A_m_-d_3_) standard.

Hana Cahová: supervised the synthesis of the m6A_m_-d_3_ standard. Liam P. Keegan: edited the manuscript.

Mary A. O’Connell: designed experiments and edited the manuscript.

Stepanka Vanacova: designed experiments, interpreted the data and wrote the manuscript.

## SUPPLEMENTARY FIGURE LEGENDS

**Figure EV1.**
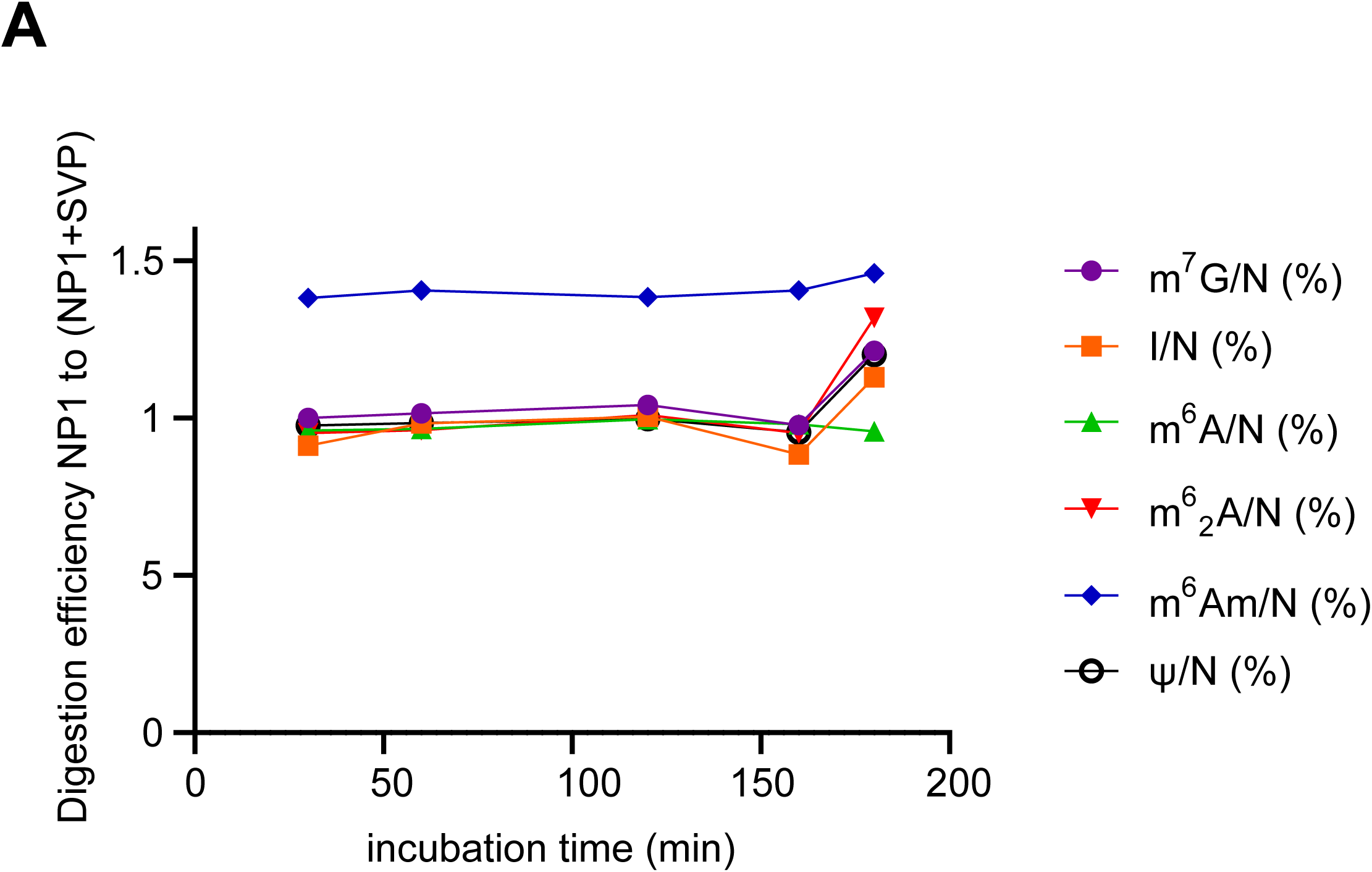
Detection of cap-linked and internal RNA modifications using the different digestion specificities of NP1 and SVP nucleases. **A.** Comparison of the cleaving activity of a mixture of NP1 and SVP (+ SVP) at different incubation times to the reaction with the NP1 enzyme only (-SVP, Incubation time 2 hours at 37°C) (n = 1). Source data are available online for this figure.

**Figure EV2.**
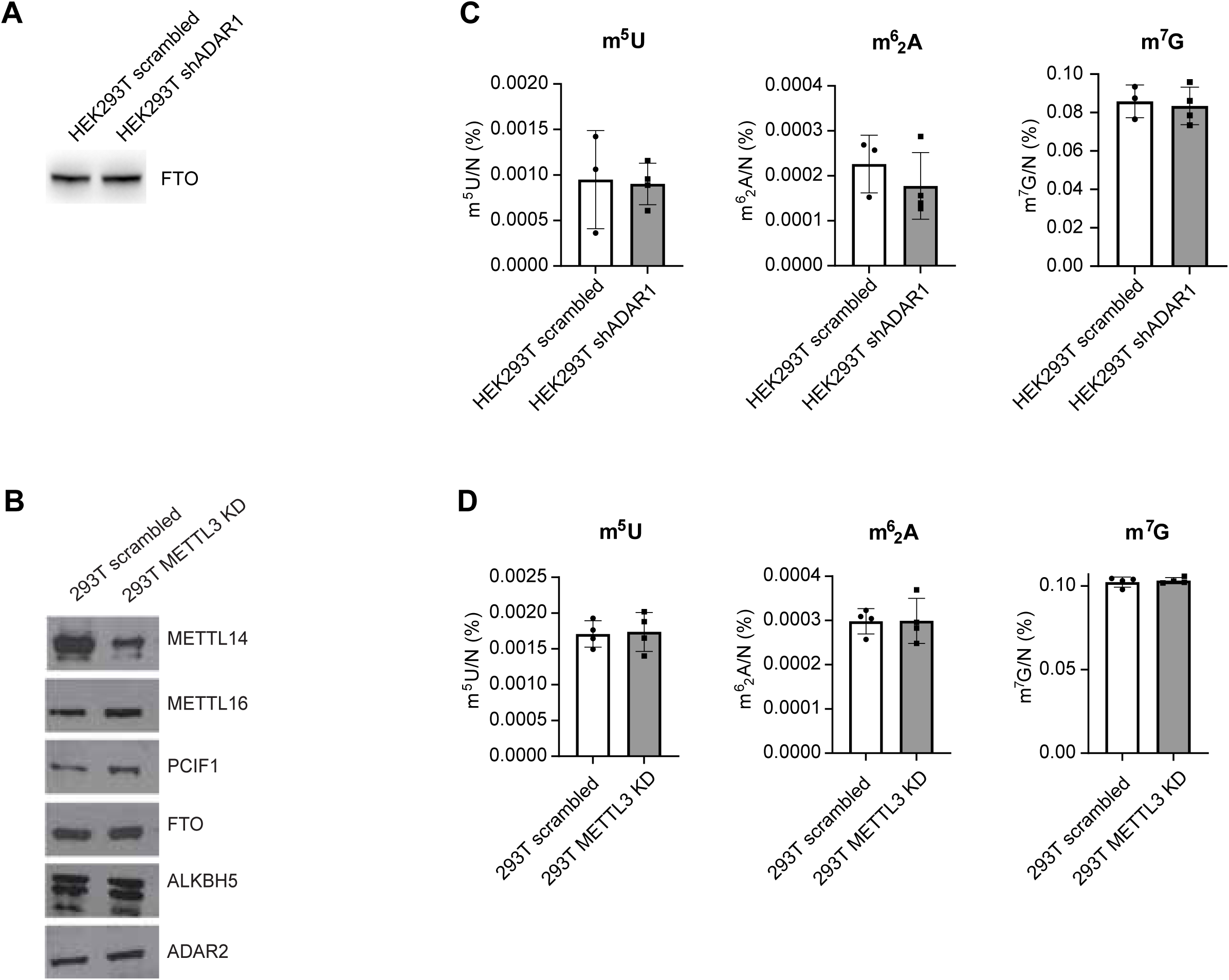
ADAR1 and METTL 3 show cross-regulation in a cell type specific manner. **A** Immunoblot analysis of protein extracts from 293T cells treated with control (scrambled) or shRNA specific for *ADAR1* mRNA. The levels of proteins were detected with specific antibodies listed in Materials and Methods. **B** Immunoblot analysis of protein expression in 293T cells transfected with control (scrambled) or *METTL3* targeting siRNAs. **C.** LC-MS/MS measurements of m^7^G and contaminating modifications originating primarily from 18S rRNA (m^6^_2_A) and tRNA (m^5^U) in poly(A) RNA after ADAR1 knockdown in HEK293T cells (n = 3 - 4). **D.** LC-MS/MS measurements of poly(A) RNA after METTL3 knockdown in 293T cells (n = 3 - 4). Data are mean ± SD, unpaired two-tailed T-test. Source data are available online for this figure.

**Figure EV3.**
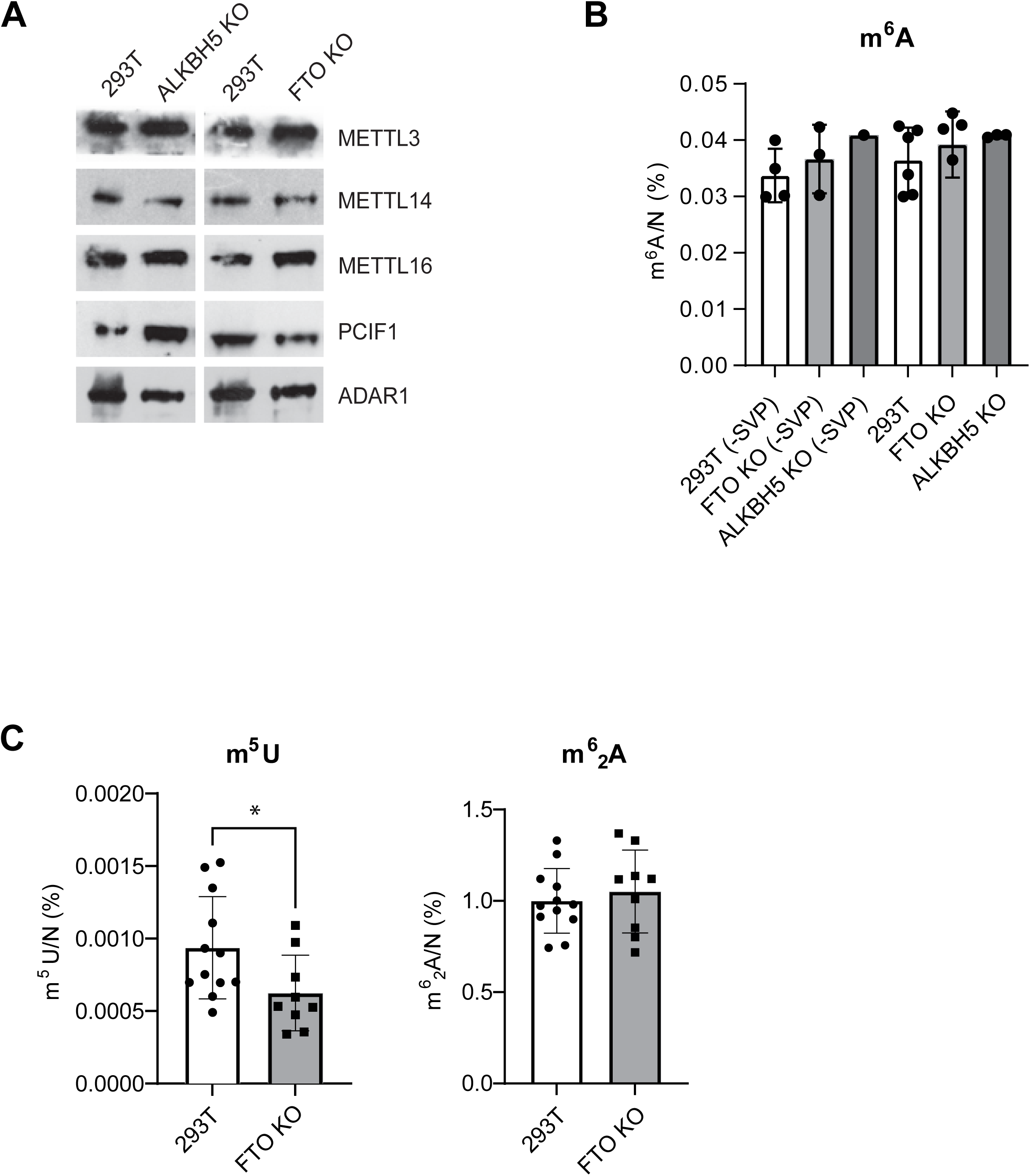
FTO primarily targets cap-associated m^6^A_m_ in noncoding RNAs. **A.** Immunoblot analysis of protein expression in 293T ALKBH5 KO and FTO KO cells. The levels of proteins were detected with specific antibodies listed in Materials and Methods. **B.** LC-MS/MS measurements and quantification of relative levels of internal (-SVP) and cap+internal m^6^A in total RNA isolated from WT, FTO KO and ALKBH5 KO 293T cells. 1 ug of total RNA was digested with NP1 and with or without SVP, to release or not the cap-linked nucleotide, respectively. In both cases, dephosphorylation with shrimp alkaline phosphatase (SAP) was performed prior to LC-MS/MS analysis. 293T (n=6); 293T (-SVP), FTO KO (n=4); FTO KO (-SVP), ALKBH5 KO (n=3); ALKBH5 KO (-SVP) (n=1) **C.** LC-MS/MS measurements of contaminating modifications originating primarily from 18S rRNA (m62A) and tRNA (m^5^U) in poly(A) RNA isolated from 293T WT (n=12) and FTO KO cells (n=9). Data are mean ± SD, One-way ANOVA, multiple comparison using the Tukey test. *P < 0.05, **P < 0.01, *** P < 0.001. The levels of nucleoside modifications were normalized to the total molar amount of canonical nucleosides N (N = C+U+G+A). Source data are available online for this figure.

**Figure EV4.**
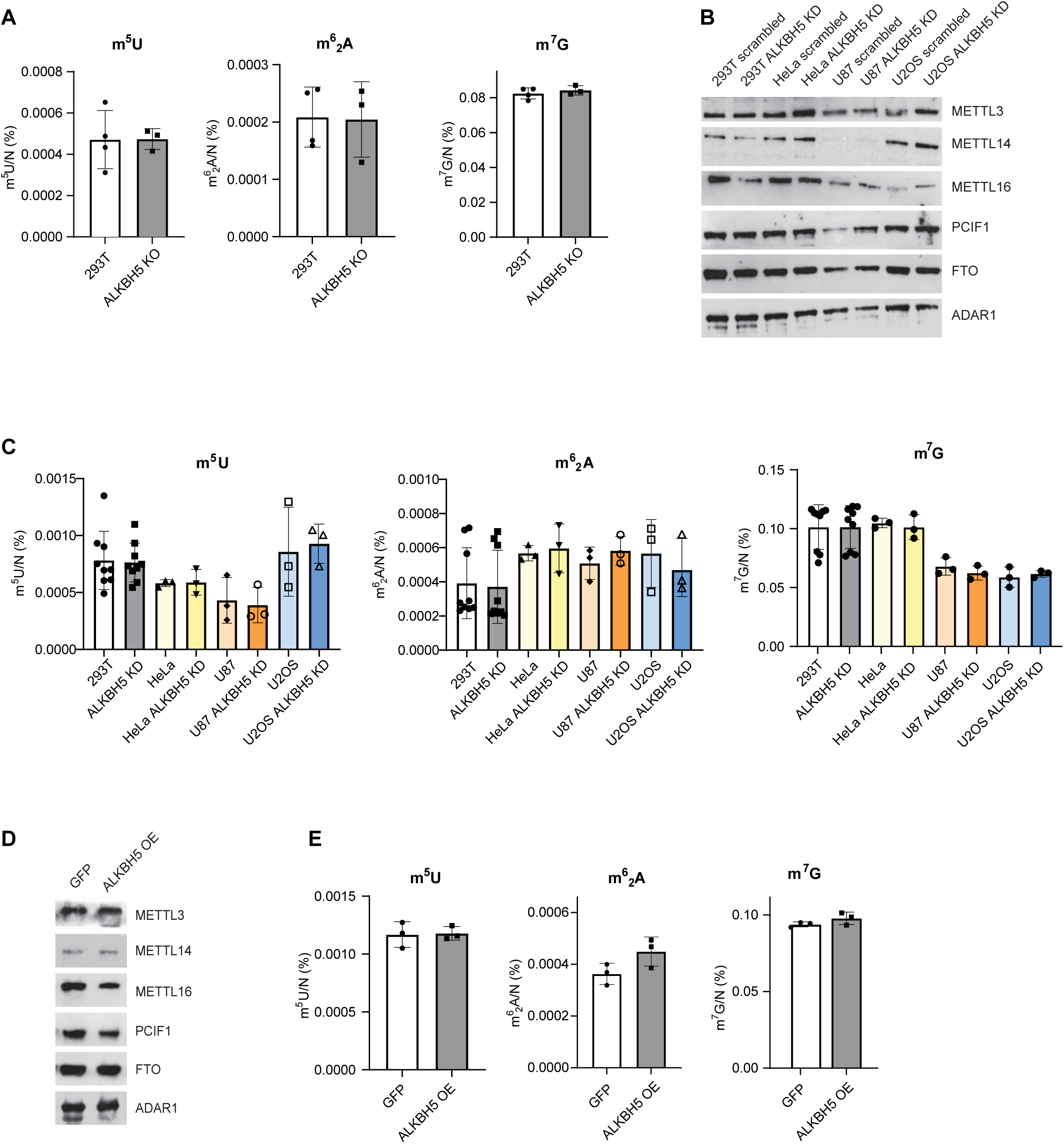
ALKBH5 depletion has a minor effect on m^6^A levels in poly(A) RNA. **A.** LC-MS/MS measurements of m^7^G and contaminating modifications originating primarily from 18S rRNA (m^6^_2_A) and tRNA (m^5^U) in poly(A) RNAs isolated from WT and ALKBH5 KO 293T cells. **B.** Immunoblot analysis of protein expression in 293T, HeLa, U87 and U2OS cells transfected with control (scrambled) or ALKBH5 targeting siRNAs (ALKBH5 KD). The levels of proteins were detected with specific antibodies listed in Materials and Methods. **C.** LC-MS/MS measurements of m^7^G and contaminating modifications originating primarily from 18S rRNA (m^6^_2_A) and tRNA (m^5^U) in poly(A) RNAs isolated from WT and ALKBH5 KD in 293T, HeLa, U87 and U2OS cells. 293T (n=9), HeLa, U87 and U2OS (n=3). **D.** Immunoblot analysis of protein expression in 293T cells overexpressing ALKBH5 (ALKBH5 OE) or GFP control. The levels of proteins were detected with specific antibodies listed in Materials and Methods. **E.** LC-MS/MS measurements of m^7^G and contaminating modifications originating primarily from 18S rRNA (m^6^_2_A) and tRNA (m^5^U) in poly(A) RNAs isolated from control (GFP) and ALKBH5 overexpressing 293T cells. Data are mean ± SD, One-way ANOVA, multiple comparison using the Tukey test (for ALKBH5 KD). Data are mean ±SD, unpaired two-tailed T-test (for ALKBH5 KO and ALKBH5 OE). *P < 0.05, **P < 0.01, *** P < 0.001. The levels of nucleoside modifications were normalized to the total molar amount of canonical nucleosides N (N = C+U+G+A). Source data are available online for this figure.

